# Di(isononyl) cyclohexane-1,2-dicarboxylate (DINCH) Alters Transcriptional Profiles, Lipid Metabolism and Behavior in Zebrafish Larvae

**DOI:** 10.1101/2021.06.11.447765

**Authors:** Noha Saad, Ceyhun Bereketoglu, Ajay Pradhan

## Abstract

Plasticizers are commonly used in different consumer goods and personal care products to provide flexibility, durability and elasticity to polymers. Due to their reported toxicity, the use of several plasticizers including phthalates has been regulated and/or banned from the market. Di(isononyl) cyclohexane-1,2-dicarboxylate (DINCH) is an alternative plasticizer that was introduced to replace toxic plasticizers. Increasing global demand and lack of information regarding toxicity and safety assessment of DINCH have raised the concern to human and animal health. Hence, in the present study, we investigated the adverse effects of DINCH (at concentrations ranging from 0.01 to 10 μM) in early developmental stages of zebrafish using different endpoints such as hatching rate, developmental abnormalities, lipid metabolism, behavior analysis and gene expression. We found that DINCH caused hatching delay in a dose-dependent manner and altered the expression of genes involved in stress response. Lipid staining using Oil Red O stain showed a slight lipid accumulation around the yolk, brain, eye and neck with increasing concentration. Genes associated with lipid metabolism such as fatty acid synthesis, β-oxidation, elongation, lipid transport were significantly altered by DINCH. Behavioral analysis of larvae demonstrated a distinct locomotor activity including distance and acceleration upon exposure to DINCH both in light and dark. Genes involved in cholesterol biosynthesis and homeostasis were also affected by DINCH indicating possible developmental neurotoxicity. The present data shows that DINCH could induce physiological and metabolic toxicity to aquatic organisms. Hence, further analyses and environmental monitoring on DINCH should be conducted to determine its safety and toxicity levels.

## 1. Introduction

Plasticizers are multifunctional chemicals used to provide flexibility, durability and elasticity to polymers by reducing their glass transition temperature, melt viscosity and elastic modulus (Bui et al., 2016; Wypych, 2004). Phthalate plasticizers are widely used in consumer goods and personal care products and comprise more than 80% of the global plasticizer market of polyvinyl chloride plastic production (Chiellini et al., 2013; Crinnion, 2010). Phthalates do not have covalent interaction with the products, hence, they leach out and contaminate the environment (Crinnion, 2010). Their high and widespread use has resulted in the detection of their metabolites in urine samples of general population from U.S., Europe and Canada (de Renzy-Martin et al., 2014; Mínguez-Alarcón et al., 2016; Saravanabhavan et al., 2013; Zota et al., 2014). Phthalates have been shown to have many adverse health effects including carcinogenesis, cardiotoxicity, hepatotoxicity, nephrotoxicity, neurotoxicity and reprotoxicity (Braun et al., 2013; Diamanti-Kandarakis et al., 2009; Dodge et al., 2015; Gore et al., 2014; Messerlian et al., 2016; Miodovnik et al., 2014). Based on the reported toxicity, several phthalates have been regulated or their use in various products is banned in Europe (Fontelles and Clarke, 2005), U.S. (CPSIA, 2008), and Canada (CCPSA, 2020). Increasing demand for safer and environmentally friendly plasticizers has led the industry to investigate and produce non-phthalate plasticizers. Several alternative plasticizers are now in the market and their production is gradually increasing (Bui et al., 2016).

Di(isononyl) cyclohexane-1,2-dicarboxylate (DINCH) was introduced in the market with a trade name of Hexamoll DINCH in 2002 as a safer alternative plasticizer (EFSA, 2007). The use of DINCH in different food contact items was approved by the European Food Safety Authority in 2007 (EFSA, 2007). It is also used in many polyvinyl chloride (PVC) products such as children toys and medical devices (EFSA, 2007). The global demand for alternative plasticizers is increasing and in parallel with this, the production and consumption rate of DINCH is on the rise (BASF, 2014). In the European market, the production volume of DINCH is more than 10000 tons per year which makes it one of the most used plasticizers (ECHA, 2021).

DINCH was found to be the most abundant non-phthalate plasticizer in Swedish preschool dust with geometric mean level of 73 μg/g dust (Larsson et al., 2017). DINCH metabolites, cyclohexane-1,2-dicarboxylic acid-monocarboxy isooctyl ester (MCOCH) and cyclohexane-1,2-dicarboxylic acid-mono (hydroxy-isononyl) ester (MHiNCH) were not detected in urine samples of adult population from Germany and USA (Schütze et al., 2014; Silva et al., 2013). However, the concentrations of cyclohexane-1,2-dicarboxylic acid monoisononyl ester (MINCH) metabolites, OH-MINCH, cx-MINCH and oxo-MINCH, increased over the years and were detected to be 2.09, 0.86 and 1.81 μg/L, respectively in urine samples of the German Specimen Bank (Schütze et al., 2014). Moreover, several other studies have also detected DINCH metabolites in urine samples (Fromme et al., 2016; Kasper-Sonnenberg et al., 2019; Mínguez-Alarcón et al., 2016; Ramos et al., 2016; Schwedler et al., 2020; Urbancova et al., 2019). The tolerable daily intake is estimated to be 1 mg/kg bw/day based on the rat data (EFSA, 2006). The oral reference dose for DINCH was estimated to be 0.7 mg/kg bw/day. This dose was based on human equivalent 10% benchmark response levels of 21 mg/kg-day for thyroid growth status observed in rats (Bhat et al., 2014).

Several *in vivo* and *in vitro* toxicological studies have analyzed possible adverse effects of DINCH and contradictory results were obtained (Campioli et al., 2015; Campioli et al., 2019; Campioli et al., 2017; David et al., 2015; EFSA, 2007; Eljezi et al., 2019; Engel et al., 2018; Nardelli et al., 2017; Vasconcelos et al., 2019). Although the *in vivo* toxicological studies of DINCH on rats have shown no effect on behavior, organ weight, serum chemistry (David et al., 2015), no evidence of reproductive toxicity or endocrine disruptive properties (EFSA, 2007), higher doses (300– 1000 mg/kg body weight/day) resulted in thyroid hyperplasia and renal toxicity (EFSA, 2007). It has been suggested that one of the metabolites, MINCH is a potent PPAR-α agonist and has a potential metabolic disrupting effect that alter fat storage in adipocytes resulting in obesity (Campioli et al., 2015). The same research group also showed that DINCH alters liver gene expression in rat at a dose of 1 mg/kg bw/day (Campioli et al., 2019), affects Leydig cell function, and impairs liver metabolic capacity in *utero* exposure of rats (Campioli et al., 2017). In another study, higher incidence of hemorrhagic testes was observed in the offspring of timed-pregnant Sprague-Dawley rats that were gavaged with 30 and 300 mg/kg/day of DINCH (Nardelli et al., 2017). In a study, it has been shown that DINCH did not have any effect on the activity of human nuclear receptors ERα, ERβ, AR, PPARα and PPARγ in HEK293 cell line, while its metabolites, M2NCH, MINCH, OH-MINCH, oxo-MINCH, and cx-MINCH, were shown to activate these receptors (Engel et al., 2018). Taken together, there is still a lack of information regarding toxicity and safety assessment of DINCH. Moreover, the available data are debatable whether DINCH is of concern to human health, hence, further research should be conducted to reveal its potential risks and toxic effects on population.

Zebrafish (*Danio rerio*) has become an important vertebrate model system for biomedical and genetic research such as development, lipid metabolism and behavior (Ho et al., 2016; Kalueff et al., 2016; Santoro and Metabolism, 2014; Seth et al., 2013). It offers several advantages including easy handling, small size, and transparent body that allows for continuous visualization of developmental changes (Bailey et al., 2013). The latter characteristic is particularly important to assess structural integrity of zebrafish as well as its functional ability that can be used to determine the impact of chemicals on behavior (Bailey et al., 2013; Fitzgerald et al., 2020). Besides, it shares a higher genetic similarity with human, as around 70% of human genes have at least one ortholog in zebrafish (Howe et al., 2013). Several important genes associated with lipid metabolism in mammals such as genes involved in fatty acid transport and acyl-CoA synthesis have been found in zebrafish (Seth et al., 2013).

The present study aimed to analyze the adverse effects of DINCH at early developmental stages of zebrafish through assessment of different endpoints such as hatching rate, developmental abnormalities, lipid metabolism, behavior and gene expression. The results reveal that DINCH has negative effects on zebrafish embryonic development and provide insights into molecular mechanisms of DINCH toxicity.

## 2. Materials and Methods

### 2.1. Chemicals

DINCH (CAS No. 166412-78-8; molecular formula: C_6_H_10_(CO_2_C_9_H_19_)_2_) was purchased from Sigma (purity ≥95 %). The physical chemical properties of DINCH are as follows: vapour pressure (mm Hg) < 0.000001 hPa, water solubility < 0.02 mg/L, partitioning coefficient noctanol/water (log Pow) = 10. The stock solutions were prepared in dimethylsulfoxide (DMSO; Sigma).

### 2.2. Zebrafish maintenance

The wild type zebrafish (ORU strain) were maintained at 25±1 °C with 14 hr light/10 hr dark cycle. *Artemia salina* and flake food (Tetrarubin) were used as food source and fish were fed twice a day. To get eggs, 4-5 couples were kept in a spawning container during the evening hours and the eggs were collected the following morning. The ethical permit for zebrafish handling was approved by the Swedish Ethical Committee (Permit 5.2.18-4065/18).

### 2.3. Exposure

The eggs were collected and then transferred in E3 medium (580 mg of NaCl, 26.6 mg of KCl, 96 mg of CaCl_2_, and 163 mg of MgCl_2_ per liter). DINCH stock solutions were added to E3 medium to obtain final assay concentrations 0.01 (0.0042 mg/L), 0.1 (0.042 mg/L), 1 (0.42 mg/L), 10 (4.2 mg/L) μM. The final assay concentration of DMSO was maintained at 0.1%. For each replicate, 30 embryos (2 hours post fertilization; hpf) were exposed in a 6 well plate (Sartstedt). Each exposure was performed in triplicate. The plates were kept at 24.0 ± 1.0 ºC. Hundred percent of exposure media was changed every alternate day.

### 2.4. Hatching delay, mortality and abnormality analysis

Hatching rates, mortality, pigmentation, morphological and developmental abnormalities were monitored and noted each day. Any malformations including spinal defects and pericardial edema were checked under the stereomicroscope (Lumar.V12 /Zeiss) connected to AxioVision 4.7.1 software (Zeiss).

### 2.5. Lipid staining

The 0.5% stock solution of Oil Red O (ORO) dye (Sigma) was prepared in 100% isopropanol and was diluted to the 0.3% working solution using 60% isopropanol. Zebrafish larvae at 6 dpf was collected and fixed using 4% paraformaldehyde for 1 hr at room temperature (RT). The larvae were then washed twice with phosphate-buffer saline (PBS) solution. Subsequently, the larvae were fixed with 60% isopropanol for 30 min and washed again with PBS. The larvae were then stained with ORO working solution for 1 hr at RT. The larvae were washed three times with 60% isopropanol. Images were taken with 4× objective using a bright field microscope (Olympus BX51).

### 2.6. Behavior analysis

The locomotor response of zebrafish larvae to a light-dark transition was investigated. On day 6, larvae were transferred to 96 well plate containing 250 µl of exposure solution and allowed to acclimatize for 4 hr. Alteration in locomotion was then analyzed using Danio Vision (Noldus). During the recording, temperature was maintained at 25 °C. For stimulation, light to dark transition was used to observe any behavioral change. The parameters consisted of light on for 10 min (5 min for acclimatization and 5 min for recording), light off for 5 min and then light on for 5 min recording.

### 2.9. RNA isolation and Quantitative real-time PCR (qPCR)

For gene expression analysis, 0.01, 0.1, 1 and 10 μM of DINCH were used. For each exposure group, 8 biological replicates were used and for each biological replicate, 4 larvae at 6 dpf were pooled in a homogenizing tube. The samples were lysed in lysis buffer (Macherey Nagel) and RNA extraction was performed using RNA extraction kit (Macherey Nagel). cDNA was prepared using the cDNA synthesis kit (Quanta Biosciences). The qPCR was carried out on thermocycler (CFX96; BioRad) using SYBR Green (PCR Biosystems). The qPCR cycles consisted of an initial denaturation at 95 °C for 2 min, 40 cycles of 95 °C for 5 s and 60 °C for 30s. The data was normalized using *elongation factor* (*eef1a1*) and ΔΔCt method was used to calculate fold change (Schmittgen and Livak, 2008). Primer sequences are listed in Table S1.

### 2.10. Statistical Analysis

To determine if control and DINCH exposed groups were significantly different, one-way analysis of variance (ANOVA) and Dunnett post-hoc test using the GraphPad Prism 8 software (GraphPad Software) were performed. For behavior analysis, Kruskal-Wallis test was used. The differences were considered significant when the p value was <0.05 (*p < 0.05; **p < 0.01; ***p ≤ 0.001, ****p<0.001).

## 3. Results

### 3.1. DINCH decreases hatching rates and induces stress

Exposure to DINCH delayed hatching rates in dose and time-dependent manner. At 80 hpf, all the doses of DINCH significantly reduced hatching rate. The hatching rate was 26.1% in the control group, while it was 7.3%, 11.7% and 7.8% for 0.01, 0.1 and 1 μM of DINCH, respectively. Meanwhile, no hatching was observed for 10 μM of DINCH at 80 hpf. The hatching rates continued to be low for 1 and 10 μM of DINCH at 96 hpf with 66.5% and 66.6%, respectively, while it was 85.0% for the control group. However, no difference was observed at 104 and 120 hpf (Fig. 1). Exposure to DINCH did not induce any significant change on mortality at analyzed DINCH concentrations (data not shown). DINCH caused slight edema and swelling in yolk region in response to 10 μM, while no other malformation was observed in response to DINCH.

**Figure 1.**
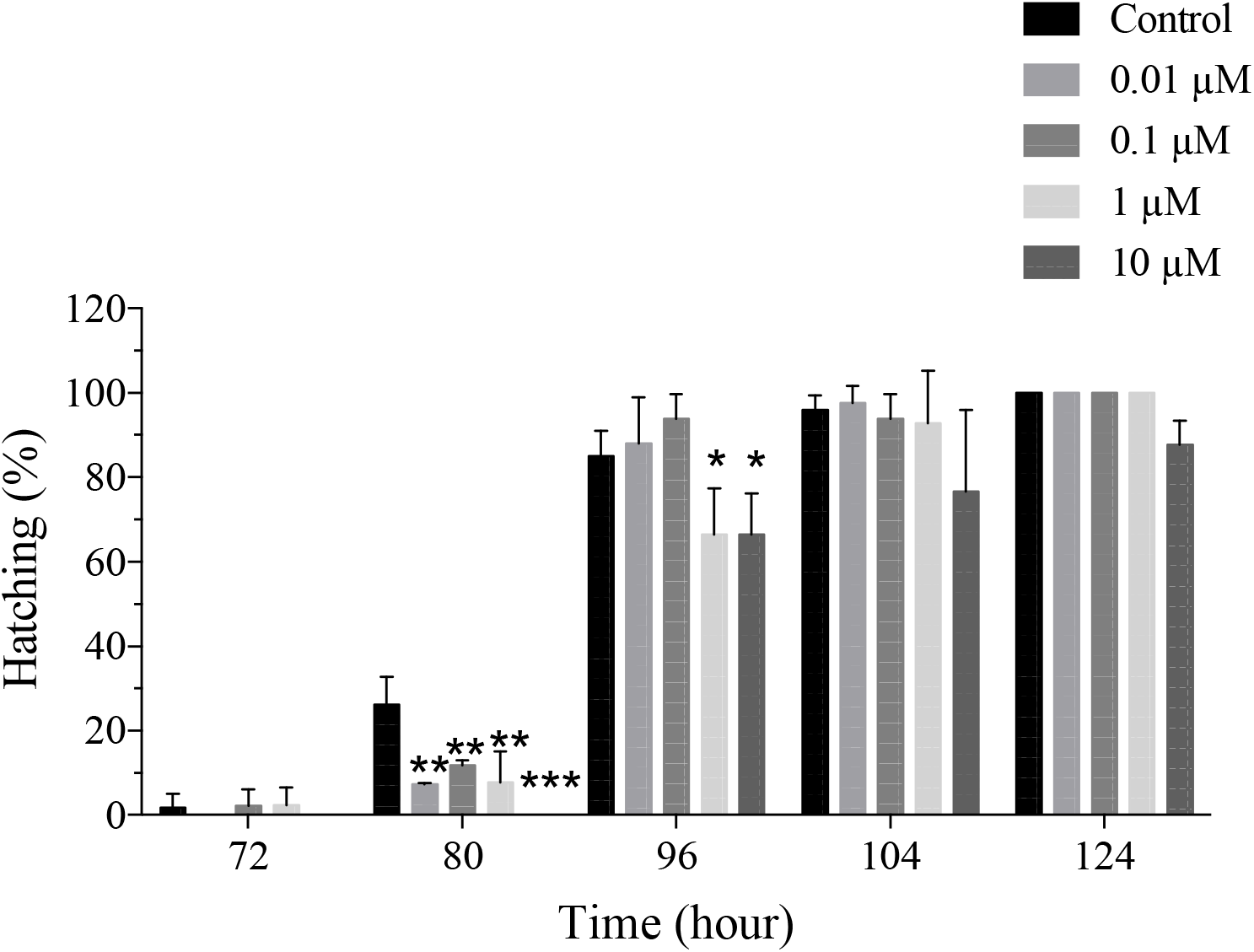
DICNH leads to hatching delay. Zebrafish eggs at 2 hpf were exposed to different concentrations of DINCH and hatching delay was recorded until 124 hpf. Statistical analysis was performed using One-way ANOVA followed by Dunnett’s post-test. Error bars represent mean±SD, n = 90.

To determine whether DINCH induces stress in zebrafish, expression of stress-related genes was analyzed by qRT-PCR upon exposure to 0.01, 0.1, 1, and 10 μM of DINCH for 120 hpf (0 to 6 days). The results showed that DINCH can induce oxidative stress response by altering the expression of various genes. Downregulation of superoxide dismutase genes, *sod1, sod2* and *sod3* was observed in response to all doses of DINCH (Fig. 2A, B and C). The *glutathione s-transferase* (*gst*) was also downregulated upon exposure to all the doses of DINCH (Fig. 2D). The gene involved in apoptosis *cytochrome c 1* (*cyc1*) was significantly upregulated by 0.01, 0.1 and 10 μM of DINCH (Fig. 2E), while *FKBP prolyl isomerase 4* (*fkbp4*) which is involved in oxidative stress was induced in response to 1 and 10 μM of DINCH (Fig. 2F). On the other hand, *catalase* (*cat*) did not show any significant change (Fig. 2G).

**Figure 2.**
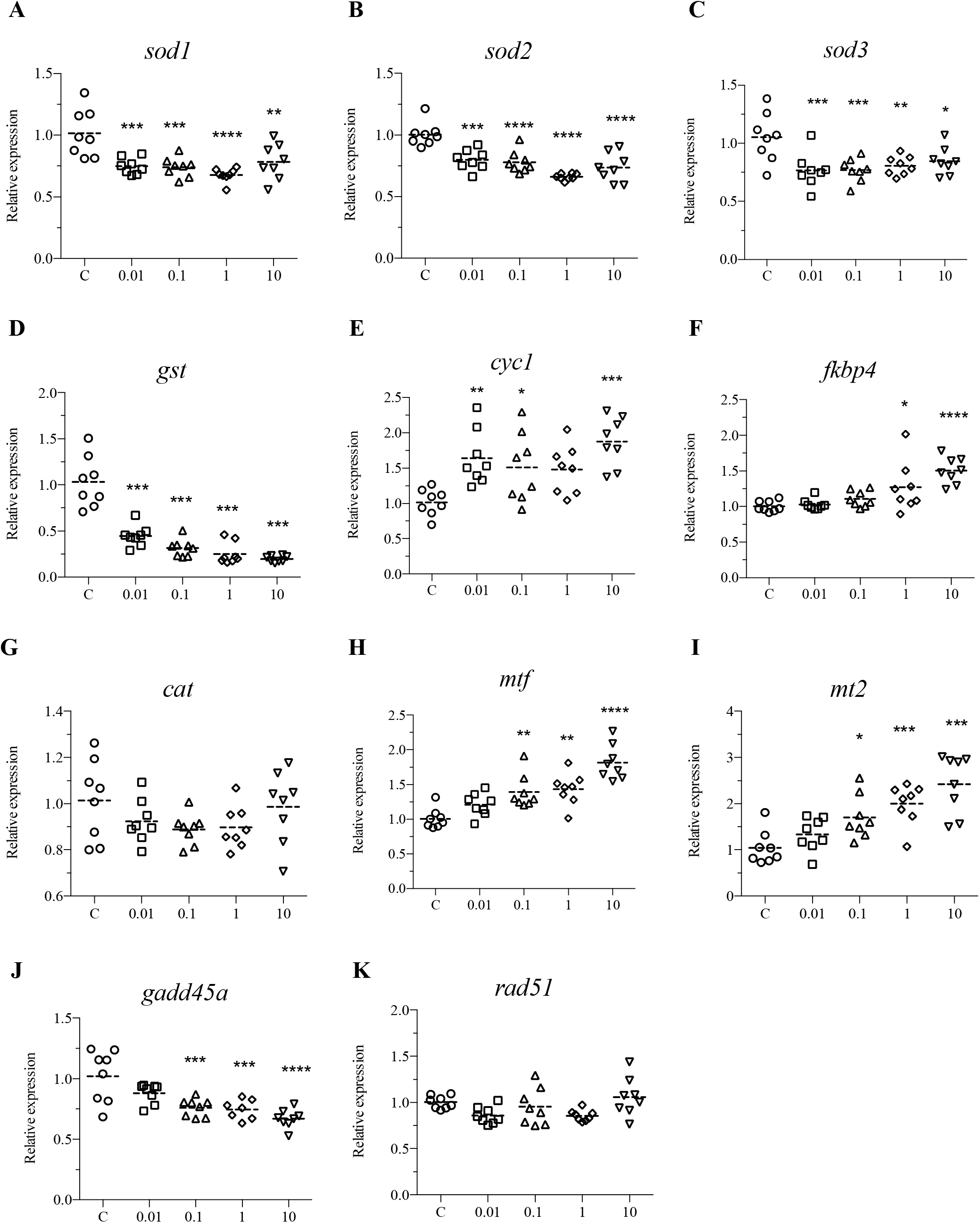
DINCH affects the expression of oxidative stress genes. Zebrafish embryos were exposed to 0.01, 0.1, 1 and 10 µM of DINCH for 6 dpf and qRT-PCR analysis was performed for stress related genes including *sod1* (A), *sod2* (B), *sod3* (C), *gst* (D), *cyc1* (E), *fkbp4* (F), *cat* (G), *mtf* (H), *mt2* (I), g*add45a* (J) and *rad51* (K). Statistical analysis was performed using One-way ANOVA followed by Dunnett’s post-test.

Genes involved in metal stress, *metal regulatory transcription factor* (*mtf*) and *metallothionein 2* (*mt2*) were significantly upregulated upon exposure to 0.1, 1 and 10 μM of DINCH (Fig. 2H and I). Of the genes involved in cell cycle and DNA damage, *growth arrest and DNA-damage-inducible, alpha* (*gadd45a*) was downregulated in response to 0.1, 1 and 10 μM of DINCH (Fig. 2J), while *RAD51 recombinase* (*rad51*) did not show change at any exposure condition (Fig. 2K).

### 3.2. DINCH alters lipid metabolism

Lipid staining was performed to determine if DINCH affects lipid metabolism in zebrafish larvae. The results showed that there was a modest increase in lipid content as the intensity and localization of ORO stain increased around the yolk region (Fig. 3). ORO staining was also localized in the brain, around the eye, neck, and heart in a dose-dependent manner (Fig. 3).

**Figure 3.**
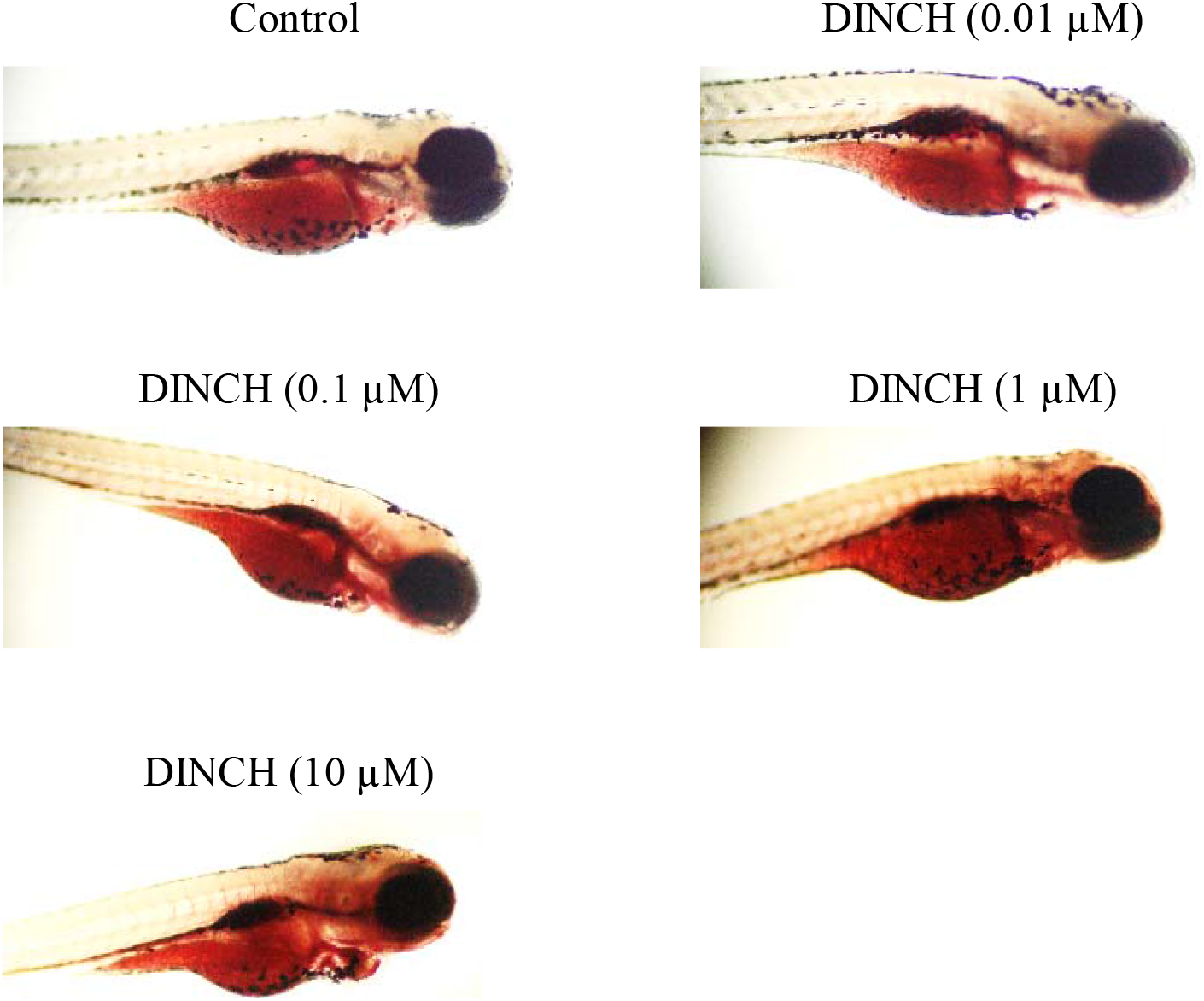
DINCH alters lipid metabolism. Zebrafish embryos were exposed to 0.01, 0.1, 1 and 10 µM of DINCH for 6 dpf and ORO staining was performed. Images were taken with 4X objective using a bright field microscope (Olympus BX51). n = 10.

To further understand the reason for lipid metabolism alteration, gene expression analysis was performed. Expression profiles showed that several genes associated with lipid metabolism were regulated upon exposure to DINCH. *sterol regulatory element binding transcription factor 1* (*srebp1*) was significantly upregulated by 0.1, 1, and 10 μM doses (Fig. 4A), while *srebp2* was only induced by 10 μM (Fig. 4B). *fatty acid synthase* (*fasn*) was upregulated in response to 0.1, 1, and 10 μM doses (Fig. 4C). Other genes including *fatty acid elongase 1* (*elovl1*) was only significantly upregulated by 10 μM (Fig. 4D), while *elovl2* was repressed by 0.01 and 1 μM of DINCH (Fig. 4E). Peroxisome proliferator activated receptor genes were also analyzed. Of these *pparg* was significantly downregulated in all the exposure groups (Fig. 4F), while *ppara* was upregulated in response to 0.1 and 10 μM doses (Fig. 4G). In addition, *pparb* was induced upon exposure to 0.1, 1, and 10 μM of DINCH (Fig. 4H).

**Figure 4.**
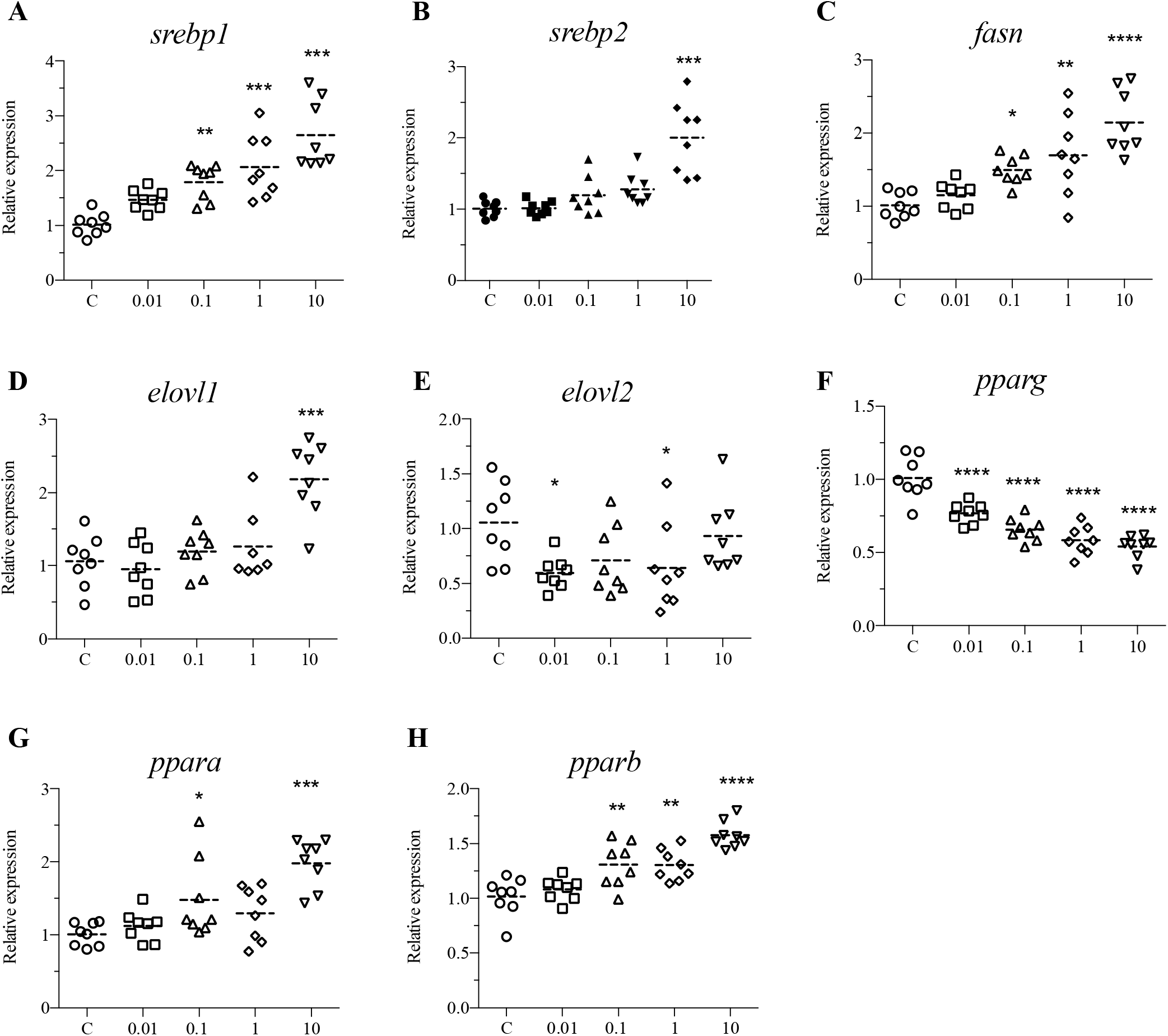
DINCH alters fatty acid synthesis, peroxisome proliferator activated receptor, and fatty acid elongation genes. Zebrafish embryos were exposed to 0.01, 0.1, 1 and 10 µM of DINCH for 6 dpf and qRT-PCR analysis was performed lipid metabolism genes including srebp1 (A), srebp2 (B), fasn (C), *srebp1* (A), *srebp2* (B), *fasn* (C), *elovl1* (D), *elovl2* (E), *pparg* (F), *ppara* (G) and *pparb* (H). Statistical analysis was performed using One-way ANOVA followed by Dunnett’s post-test.

Apolipoprotein genes were also significantly altered. Of these, *apoeb* and *apoa4* were downregulated by all exposure groups (Fig. 5A and B), while a*poa1* was downregulated by 0.1, 1, and 10 μM doses (Fig. 5C). *StAR-related lipid transfer domain containing 4* (*star4d*) which is involved in cholesterol transport was upregulated only by 10 μM DINCH (Fig. 5D). Other genes including *low density lipoprotein receptor* (*ldlr*) were significantly downregulated by all the exposure concentrations (Fig. 5E), while *lipase C* (*lipc*) was induced by 1 and 10 μM of DINCH (Fig. 5F). Genes involved in fatty acid cleavage including *phospholipase B* (*plb*) were downregulated by 0.01, 0.1 and 10 μM of DINCH (Fig. 5G), while *lipoprotein lipase* (*lpl*) did not show significant change at any exposure condition (Fig. 5H).

**Figure 5.**
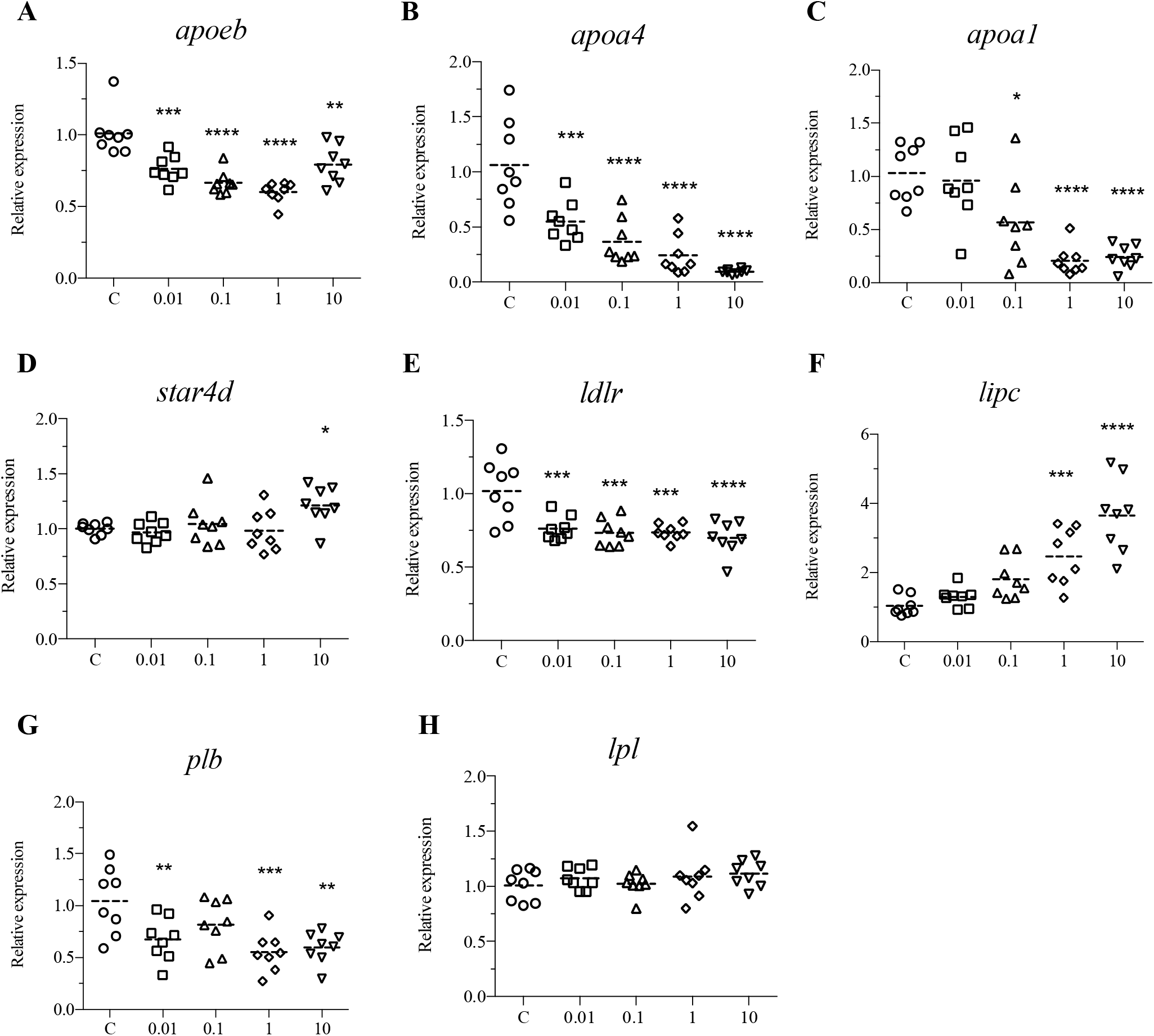
DINCH alters lipid transport genes. Zebrafish embryos were exposed to 0.01, 0.1, 1 and 10 µM of DINCH for 6 dpf and qRT-PCR analysis was performed for lipid transport and processing including *apoeb* (A), *apoa4* (B), *apoa1* (C), *star4d* (D), *ldlr* (E), *lipc* (F) *plb* (G) and *lpl* (H). Statistical analysis was performed using One-way ANOVA followed by Dunnett’s post-test.

### 3.3. DINCH alters behavior

The light-dark locomotor activity test was performed for 6 dpf zebrafish larvae in response to 0.01, 0.1,1 and 10 μM DINCH. The distance moved did not differ in any exposure groups and the control during the first and second stimulations. There was also no difference when the light was switched off (Fig. 6A, B and C). However, the distance moved by the larvae increased in all exposure concentrations during the light off period compared to first stimulation. Besides, the distance moved by the larvae came back to the level of first stimulation in the second stimulation period (Fig. 6A, B, C). During the first stimulation, 1 and 10 μM of DINCH showed increased acceleration (Fig. 6D), while in the dark cycle, no change was observed (Fig. 6E). The 0.1 and 1 μM treated group showed increased acceleration while 10 μM DINCH did not show any difference when the light was switched on again (Fig. 6F). Interestingly, the acceleration was found to be significantly lower during the first recording compared to light off and second stimulation (Fig. 6D, E, and F).

**Figure 6.**
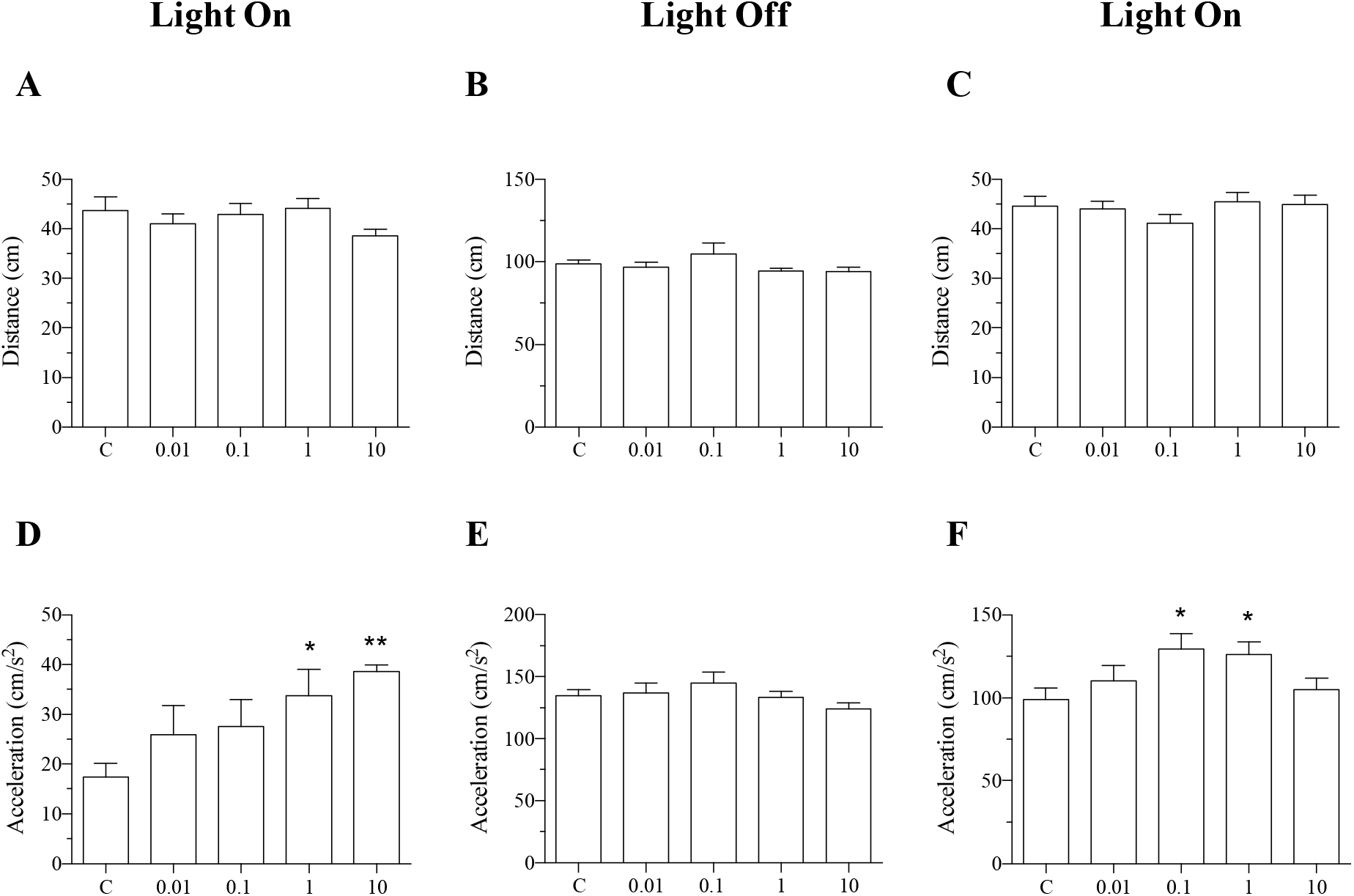
DINCH alters behavior in larvae. Zebrafish embryos were exposed to 0.01, 0.1, 1 and 10 µM of DINCH for 6 dpf and behavior analysis was performed. Distance and acceleration were recorded during light on (A and D), Light off (B and E) and light on (C and F). Statistical analysis was performed using One-way ANOVA followed by Dunnett’s post-test. n = 48.

Since behavior was altered following DINCH exposure, the expression of genes involved in behavior were also investigated. *proprotein convertase subtilisin/kexin type 9* (*pcsk9*) was significantly upregulated by all the doses (Fig. 7A), while *doublesex and mab-3 related transcription factor 3A* (*dmrt3a*) was induced in response to 0.1, 1, and 10 μM of DINCH (Fig. 7B). Meanwhile and 3-hydroxy-3-methylglutaryl-coa synthase 1 (*hmgcs1*) were upregulated by only 10 μM of DINCH (Fig. 7C). In addition, myelin basic protein a (*mbpa*) was upregulated upon exposure to 1 and 10 μM of DINCH (Fig. 7D), while *7-dehydrocholesterol reductase* (*dhcr7*) was significantly downregulated by all the doses (Fig. 7E). However, *cfos* did not show change at any exposure condition (Fig. 7F).

**Figure 7.**
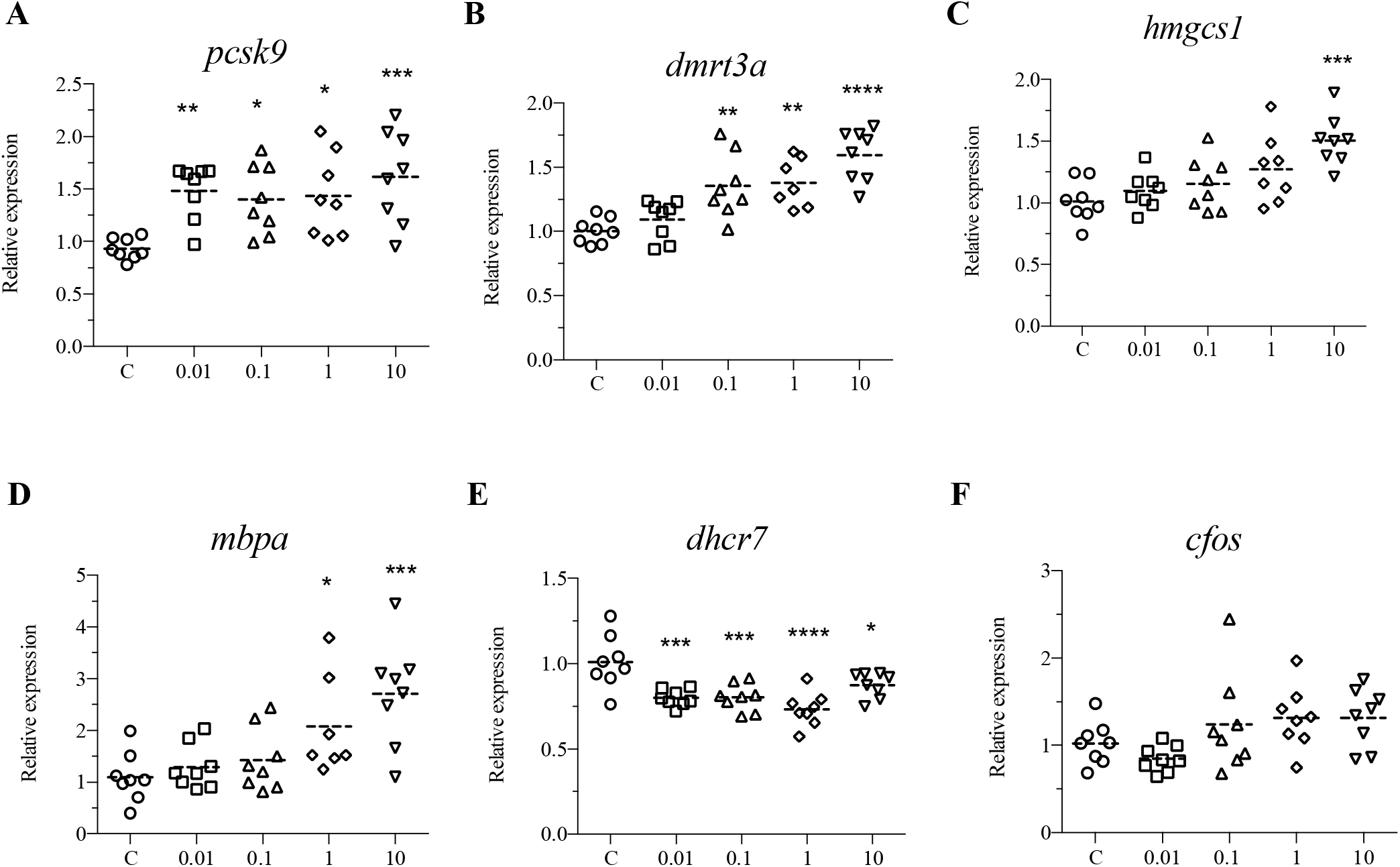
DINCH affects behavior, cholesterol biosynthesis and homeostasis genes. Zebrafish embryos were exposed to 0.01, 0.1, 1 and 10 µM of DINCH for 6 dpf and qRT-PCR analysis was performed for genes involved in behavior including *pcsk9* (A), *dmrt3a* (B), *hmgcs1* (C), *mbpa* (D), *dhcr7* (E) and *cfos* (F). Statistical analysis was performed using One-way ANOVA followed by Dunnett’s post-test.

## 4. Discussion

We observed that DINCH delayed hatching rates in dose and time-dependent manner. Although there is no study examining the effect of DINCH on zebrafish hatching, several studies reported delayed hatching in response to other plasticizers including di-n-butyl phthalate (DBP), di-(2-ethylhexyl) phthalate (DEHP), and acetyl tributyl citrate (ATBC) (Hu et al., 2020; Muhammad et al., 2018). No significant change in mortality was observed in the present study. Consistent with our observation, no mortality has been determined on rats following intravenous administration of DINCH up to 300 mg/kg body weight/day (David et al., 2015). Several *in vitro* investigations have also addressed the effects of DINCH on cell viability and demonstrated contradictory results. (Boisvert et al., 2016; Nardelli et al., 2015).

Oxidative stress is a key indicator in environmental and toxicological risk assessment. We observed downregulation of *sod1, sod2* and *sod3* by all the doses of DINCH. It has previously been indicated that these genes were affected by several plasticizers such as DEHP, dimethyl phthalate (DMP), and butyl benzyl phthalate (BBP) (Cong et al., 2020; Yang et al., 2018; Zhang et al., 2014). It has also been shown that SOD1 enzyme activity was overexpressed in response to subacute exposure to DINCH at post-natal day 21 of dams, while it was repressed upon *in utero* exposure to DINCH at post-natal day 60 in a dose-dependent manner (Campioli et al., 2019). We also observed an increased expression of *cyc1* and *fkbp4* upon exposure to DINCH. Of these, *cyc1* is involved in mitochondrial oxidative phosphorylation (Chen et al., 2015), while *fkbp4* encodes an immunophilin protein that may have a protective role against oxidative stress (Gallo et al., 2011). Taken together, we suggest that altered expression of the above-mentioned genes in response to DINCH may result in ROS generation and subsequently cause oxidative stress in zebrafish.

Metallothioneins are proteins that can bind metals and implicated in several biological processes including metal homeostasis and regulation of oxidative stress (Chen et al., 2004; Dorts et al., 2016; Ruttkay-Nedecky et al., 2013). It has been previously demonstrated that the expression of *Mt1a, Mt2a*, and *Mt1m* was significantly induced in postnatal day 60 rats treated *in utero* with 1 and 100 mg/kg/day of DINCH (Campioli et al., 2019). We observed induced expression of two genes associated with metallothionein pathway, *mtf* and *mt2* in response to 0.1, 1 and 10 μM of DINCH. This result supports that DINCH may cause oxidative stress and metallothionein related genes were overexpressed to overcome the stress. We have also analyzed the genes involved in cell cycle and DNA damage. Although the expression of *rad51* did not show any change, we found significant downregulation of *gadd45a* that promotes cell cycle arrest and DNA excision repair (Salvador et al., 2013). Altogether, these findings suggest that DINCH exposure induces oxidative stress which may result in malfunctioning of cell cycle progression, DNA damage and hatching delay.

Lipids are key molecules that are involved in signaling, membrane composition, and energy production (Anderson et al., 2011). It has been indicated that defects in lipid metabolism can cause several disorders such as obesity, diabetes, and atherosclerosis (Joffe et al., 2001; McNeely et al., 2001; Watanabe et al., 2008). The yolk consists of the yolk syncytial layer (YSL) that is a lipid-rich structure. The YSL aids in release of fatty acids and synthesis of lipoproteins which transport lipids to the embryo to support larval growth (Anderson et al., 2011; Otis et al., 2015). In the present study, we observed a modest increase in lipid content around the yolk and we speculated that lipid transport becomes less active following DINCH exposure. In a previous study, it has been indicated that DINCH did not show any difference in lipid accumulation compared to control, while its primary metabolite resulted in lipid accumulation in preadipocytes from epididymal adipose tissue of rats (Campioli et al., 2015).

To further confirm these findings, we performed qRT-PCR for genes in certain lipid metabolism pathways including fatty acid synthesis, PPAR signaling pathway, and lipoprotein transport. SREBPs are transcription factors that regulate the expression of genes involved in uptake and synthesis of cholesterol and fatty acids (Shimano, 2000). We observed an induced expression of *srebp1* and *srebp2*. The downstream gene of Srebp, *fasn* was also significantly induced upon exposure to 0.1, 1, and 10 μM of DINCH. *fasn* converts acetyl-CoA and malonyl-CoA into palmitate which is then esterified into triacylglycerides and stored in adipose tissue (Tian et al., 2013). From this, we can assume that exposure to DINCH leads to fatty acid synthesis in zebrafish. We also analyzed the expression of *elovl1* and *elovl2* which are key genes in the synthesis of long-chain mono and polyunsaturated fatty acids (Ayisi et al., 2018) and overexpression of these genes results in high availability of these fatty acids which provide an adaptation to chemical exposure. Consistent with our observations, a previous study in rats showed that the expression of several genes involved in lipid metabolism were altered in post-natal day 3 and 60 treated in utero with 1 and 100 mg/kg/day of DINCH (Campioli et al., 2019).

PPARs are involved in lipid metabolism by regulating transcription of several genes (la Cour Poulsen et al., 2012). *ppara* and *pparb* are responsible for regulating fatty acid β-oxidation (Varga et al., 2011), while *pparg* is associated with lipid storage and adipogenesis (Francis et al., 2003). We observed an upregulation of *ppara1* and *pparb* while, *pparg* was significantly downregulated. In contrast to our finding, it has been previously shown that treatment with DINCH increased the expression of *Pparg2* (Campioli et al., 2015). In the same study, one of DINCH metabolites, MINCH was shown to be PPARα agonist at 50 μM or above concentrations (Campioli et al., 2015). On the other hand, in an *in vitro* study, it has been indicated that although DINCH did not alter the reporter gene system, its metabolites induced PPARα-and PPARγ-dependent reporter gene activities in a concentration-dependent manner (Engel et al., 2018). In another study, it has been demonstrated that DINCH-treated post-natal day 21 dams showed a residual PPAR-α overexpression (Campioli et al., 2019).

Lipoproteins play a role in transporting lipid molecules to different tissues in fish (Tocher, 2003). Analysis of lipoprotein genes in our study indicated that DINCH can also affect lipid uptake and transport in zebrafish larvae. We noted a decreased expression of *apoeb* and *apoa4* by all exposure groups, while a*poa1* was repressed upon exposure to 0.1, 1, and 10 μM of DINCH. Apolipoproteins are involved in lipid transport to specific tissues through specific binding to lipoprotein receptors (Ayisi et al., 2018). We also observed downregulation of *ldlr* in response to all exposure concentrations and upregulation of *lipc* by 1 and 10 μM of DINCH. *ldlr* is important for delivering essential lipids to maintain cellular functions and concentration of cholesterol-rich lipoproteins in the circulation (Willnow, 1999), while *lipc* is associated with triglyceride hydrolysis and high-density lipoprotein cholesterol function (Neale et al., 2010). Besides, we observed a significantly repressed expression of *plb* gene that is involved in fatty acid cleavage. Our data suggest that lipid transport becomes less active after DINCH exposure and to overcome this, the genes for lipid uptake are upregulated.

Behavior is a reflection of multifactorial interactions in organisms and significant changes in behavior in response to environmental exposures can be an indicator of adverse effects (Anderson et al., 2004). In order to evaluate whether DINCH alters neuronal signaling, we conducted behavioral assay in zebrafish larvae. The altered swimming behavior following DINCH exposure suggests that DINCH may negatively regulate motor activity and result in altered swimming behavior because of neurotoxicity. Plasticizers including DEHP and di-butyl phthalate (DBP) have been shown to alter locomotor activity in zebrafish larvae by affecting spine and skeletal system development (Qian et al., 2020).

To understand the altered behavioral activity, we analyzed the expression of genes associated with behavior. Cholesterol plays a crucial role in maintaining neuronal physiology. It has been indicated that the level of cholesterol in brain is critical for proper brain function and defect in brain cholesterol metabolism is associated with different nervous system problems, including Alzheimer’s disease, Huntington’s, disease and Parkinson’s diseases (Zhang and Liu, 2015). We observed significantly altered expression of several genes involved in cholesterol biosynthesis and homeostasis such as *pcsk9, hgmcs1* and *dhcr7* (Korade et al., 2013; Mathews et al., 2014; O’Connell and Lohoff, 2020). Cholesterol plays an essential role in development of myelin in central and peripheral nervous systems (Saher et al., 2011). In line with this, we found an upregulation of *mbpa* that encodes myelin basic protein. Expression of *dmrt3a* is associated with spinal cord development and fate specification of dorsal interneurons 6 that coordinates locomotion in animals (Andersson et al., 2012). In a recent study, it was demonstrated that *dmrt3a*-knockout zebrafish has altered behavior with decreased movement, and acceleration (Del Pozo et al., 2020). We observed an increase in acceleration and parallel with this, significantly induced expression of *dmrt3a*. Altogether, this suggests that DINCH alters larval behavior by altering key genes involved in locomotion and neural function. Analysis of other genes could provide further insights into the mechanisms of behavior toxicity.

We further performed a pathway analysis and prepared a signaling pathway for the analyzed genes (Figure 8). We found that DINCH exposure particularly affected lipid metabolism by inducing fatty acid synthesis which subsequently resulted in β-oxidation and this leads to stress. Lipid transport in zebrafish was also decreased in response to DINCH and this could lead to altered behavior. To overcome this stress, Zebrafish significantly induced genes involved in cholesterol biosynthesis and homeostasis. Upregulation of these genes may also explain the accumulation of ORO staining around the brain and eye regions.

**Figure 8.**
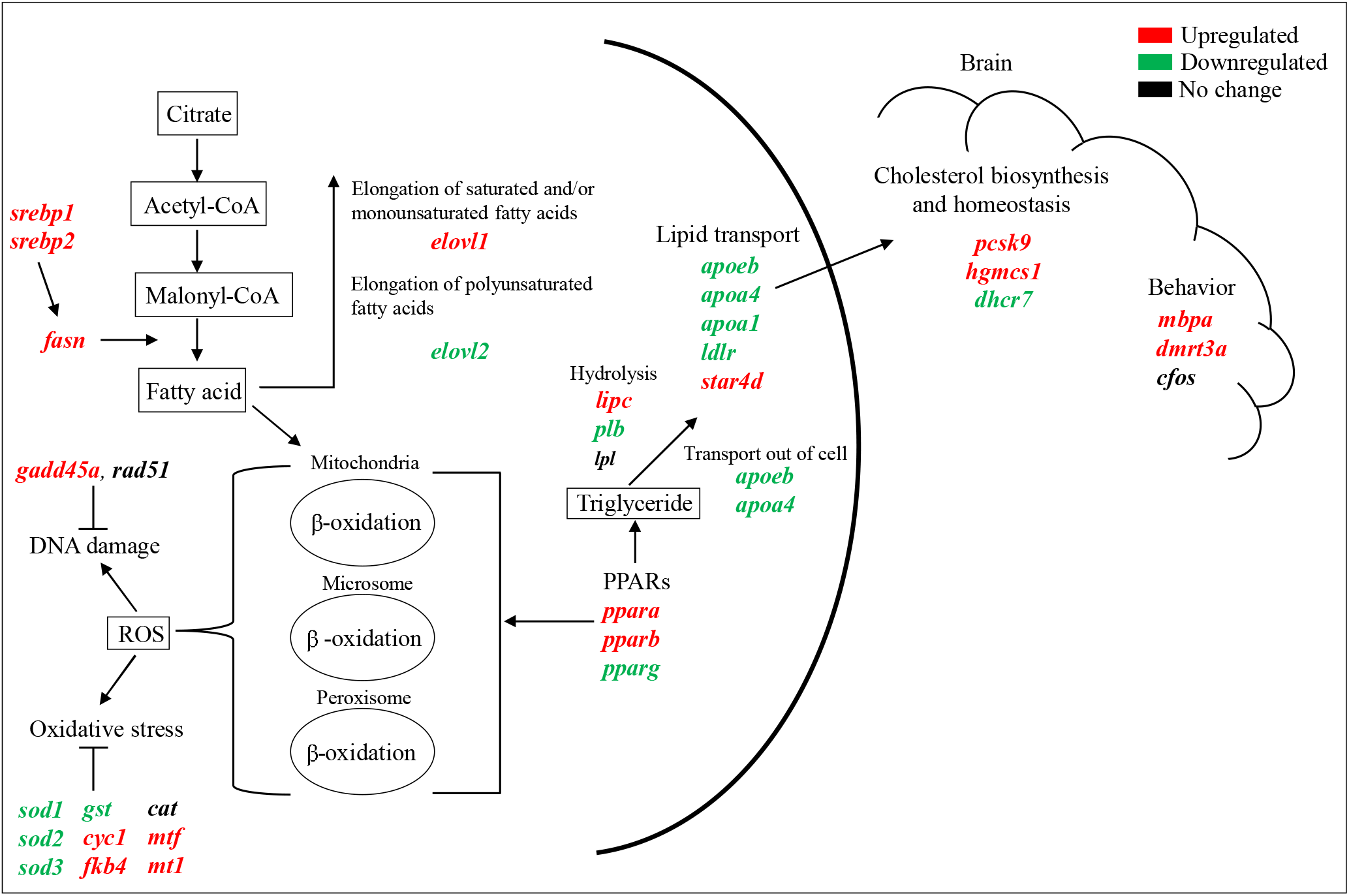
Overview of the differentially expressed genes in response to DINCH. The schematic diagram shows the regulation and involvement of genes in different signaling mechanisms. Gene involved in lipid synthesis, elongation, hydrolysis and transport were altered in larvae. The stress response and DNA repair genes were also regulated following DINCH exposure. The altered lipid metabolism and transport could also be responsible for the observed cholesterol biosynthesis/homeostasis and brain function alteration in the larvae. The upregulated genes are indicated in red color, the downregulated genes are indicated in green color and the genes that were not regulated are indicated in black color.

## 5. Conclusion

DINCH is one of the alternative plasticizers whose use is increasing in the European market. There is limited information on animal and human exposure to DINCH and the available data is conflicting as some studies have suggested that DINCH is not toxic and it cannot regulate lipid metabolism like the other plasticizers. We showed that DINCH can alter lipid metabolism and stress response genes in zebrafish larvae. Our data suggest that DINCH can have detrimental effects on aquatic organisms and higher eukaryotes. Taken together, this study indicates that DINCH alters lipid metabolism in zebrafish larvae and this alteration could also lead to changes in other physiological processes including brain functions. Further analysis on mammalian species will help to understand the risk factors associated with DINCH exposure to humans. In conclusion, DINCH is not completely safe, hence, its use, presence in the environment and negative effects on humans and animals should be carefully monitored.

## Acknowledgements

This study was financed by Knowledge Foundation Sweden, Helge Ax:son Johnsson Foundation, Längmanska Culture Foundation, Magnus Bergvalls Foundation, Örebro University, the Scientific and Technological Research Council of Turkey (TÜBİTAK, Grant No: 120Z748), and Iskenderun Technical University.

